# Lowering CD40L expression in Murine Lupus Results in an Increase in Disease Indicators in Female but not Male B6 mice

**DOI:** 10.1101/2025.02.22.639644

**Authors:** Diego Prado De Maio, Sandra Tetteh, Christopher Ayers, Michelle Brown, Lori R. Covey

## Abstract

Our lab has previously described a mouse model (CD40LΔ5) that produces only 60% of WT CD40L due to a targeted deletion in an RNA binding site within the CD40L message. The CD40LΔ5 mutation, which destabilizes CD40L mRNA during T cell activation, causes disrupted germinal center (GC) formation leading to reduced levels of memory B cells and switched antibodies. In this study, we used our model of limited CD40L expression to investigate its effect on Systemic Lupus Erythematosus (SLE or lupus). Two mouse models of SLE were assessed: The first, termed PIL, used pristane to induce disease over a six-month period and the second utilized a chronic graft-versus-host disease (bm12-cGVHD) that resembles lupus based on multiple parameters and allowed us to monitor the early events in disease development. Importantly, we found that in both model systems female mice expressing the CD40LΔ5 mutation showed a consistent *increase* in elevated antibody secreting cells and autoantibody titers. Also, PIL female CD40LΔ5 mice had higher levels of immunocomplex deposition in the kidney compared to all other cohorts. In the bm12-cGVHD model, cellular increases in GC cells along with an altered cytokine profile of donor CD4+ T cells and host dendritic cells (DCs) reflected a significant skewing of CD40LΔ5 female CD4 T cells towards a Th2 phenotype. Overall, our results support a more nuanced role for CD40L in lupus than previously described and suggest a sex-determined threshold of CD40L-CD40 signaling that demarcates an interface between protection and exacerbation at the very early steps of disease progression.

## Introduction

The dyad of CD40-CD40L is a fundamental co-stimulatory pathway establishing and orchestrating both humoral and cell-mediated immunity (1). Interactions between CD40L expressed on activated CD4+ T cells and CD40 on B cells and dendritic cells (DCs) as well as other antigen-presenting cells (APCs), catalyze essential functional immunological processes including antibody production, memory development and the magnitude and limit of local and systemic inflammation (2). In T-dependent (TD) immune responses, early engagement of CD40L with antigen (Ag)-selected B cells leads to the rapid proliferation and differentiation into short-lived extrafollicular plasmablasts in addition to cells that seed B cell follicles, which ultimately develop into germinal centers (GCs) (3–6). Importantly, *enhanced* CD40 signaling at this early stage of the response results in a preferential shift towards plasmablast differentiation and limits B cell entrance into the follicles(1, 7–9). CD4^+^CXCR5^+^ PD-1^+^ T cells are early T follicular helper cells (Tfh) that are critical for establishing the structure and function of the GCs by engaging with B cells at the border of the B cell follicles (10–13). This determinative process is driven, in part, by presenting sufficient levels of CD40L to Ag-selected B cells, thus allowing the B cell clones to proliferate, undergo affinity maturation and selection within the light and dark zones of the GC structure (14–18).

CD40 signaling is also critical for the maturation of DC and the subsequent expression of cytokines including IL-12 and IL-6 (2, 19, 20). The production of IL-12 by matured DC is essential for the expression of IFN-γ by CD4+ T cells resulting in a dominant Th1 response (21, 22). Therefore, CD40 signaling is fundamental for DC maturation as defined by the acquisition of key characteristics required to induce T-cell activation and polarization (23).

The importance of context-specific bi-directional CD40-CD40L signaling between APCs and T cells to collectively shape the quality of a TD immune response is evident. However, an understanding of pathways that regulate the expression of CD40L throughout the course of this response is not fully defined. We have recently shown that a pathway of posttranscriptional regulation of CD40L mRNA is critical for the survival and differentiation of GC B cells. This pathway is centered on an RNA stability element located in the 3’untranslated region (UTR) which was shown to regulate the turnover of the CD40L transcript in an activation-dependent manner (24). The CD40L element binds a polypyrimidine tract-binding protein-1 (PTBP1) complex that confers stability on the transcript at late times of activation (25). We engineered mice (termed CD40LΔ5) that lacked the CD40L stability element to understand the biological importance of this pathway during a humoral immune response. Notably, following challenge with NP-KLH or SRBCs, the expression of CD40L remained at approximately 60% of WT levels on Tfh cells from CD40LΔ5 mice, and this decreased expression of CD40L coincided with disorganized GCs, decreased levels of primary and secondary antibodies and impaired memory B cell development. Importantly, the CD40LΔ5 mutation resulted in a failure of B cells to proliferate and survive within the GC(26).

The impact of reduced CD40L expression on humoral immunity in a TD response led us to consider how this regulatory pathway may affect disease outcomes in systemic lupus erythematosus (SLE), an autoimmune disease driven by the high production of class-switched autoreactive antibodies to nuclear antigens. In SLE, B cell responses can be driven by the GC reaction and Tfh cells (27–31) as well as by B cell differentiation at extrafollicular (EF) sites (32–35). CD40L expression has been shown to be upregulated on both CD4+ and CD8+ cells from SLE patients (36, 37). Also, findings from transgenic mice that ectopically express CD40L expression on monocytes and B cells suggested that enhanced CD40L expression is sufficient to cause SLE (38–40). Notably, CD40 activation on B cells contributes to a feed-forward loop by upregulating CD40L surface expression on CD4+ T cells, which in turn engages CD40 on B cells (41).

Early work by several groups revealed the potential of CD40L as a highly effective therapeutic target for the treatment of autoimmune disease based on its ability to directly regulate inflammation. Ablating CD40L through targeted deletion of the gene or with antibodies against CD40L or CD40 resulted in better disease outcomes in mouse models of autoimmunity including lupus (42). In these studies, blocking CD40L-CD40 interactions in vivo interfered with the induction and maintenance of activity, the production of autoreactive antibodies, and immune complex deposition in kidneys resulting in a complete absence of end-stage renal disease(43–46). These results, in conjunction with the fact that the CD40L-CD40 pathway is a driver of SLE at different stages of disease progression, are central to the basis of developing therapeutics that focus on disrupting the development of pathogenic antibodies through this pathway (43, 47, 48). Although early results of CD40L-based therapeutics showed promise in animal models, unpredicted thrombotic events due to off-target binding on activated platelets effectively halted the development of these reagents (49, 50). Importantly, efforts to optimize the potential of CD40L as a target have continued into second generation compounds that directly address the off-target effects of the early trials (45, 51, 52). In this context, better understanding of how CD40L expression levels impact disease pathogenesis is both required and scientifically significant.

In this study we utilized two separate models of SLE to study the effect of limiting CD40L on disease development in C57B/6 (B6) mice (53–57). The pristane-induced lupus (PIL) and the bm12 chronic graft-versus-host disease (bm12-cGVHD) models allowed us to study SLE in both the early and the late stages of disease; especially prior to and following development of kidney pathogenesis. Metrics that were used to analyze the development of disease included (i) the presence of autoreactive antibodies against dsDNA and ribonuclear complexes, (ii) the deposition of immunocomplexes in the kidney glomeruli, (iii) the expansion of GC B and Tfh cells, and (iv) the differentiation of B cells into CD138+ antibody-secreting cells (ASCs) (56, 58, 59). We anticipated that mice with lowered CD40L would exhibit less severe symptoms. However, we observed only a decrease in symptoms in male CD40LΔ5 mice treated with pristane. In contrast, a consistent increase was observed in disease metrics in female CD40LΔ5 mice relative to WT male, WT female mice and CD40LΔ5 male mice. To explore these findings further, the CD40 signaling pathway in B cells was analyzed, as well as the CD4+ T cell cytokine milieu. Although we found no sex-based differences in B cell signaling, there was a clear gender-specific difference in the expression of cytokines by both CD4+ T cells and DCs suggesting a role for optimal CD40L in dampening an autoimmune response at early stages of disease. Overall, our results reveal a complex relationship between CD40L levels and SLE that is sex-determined and distinctly impacts the development of autoreactive antibodies and the severity of disease.

## Materials and Methods

### Animals

The generation of the CD40LΔ5 knock-in mice has been previously described (26). Briefly, a targeting construct containing a deleted region of 175 bp in the 3’UTR, which included the stability regions of the CD40L transcript, was introduced into B6 mice by targeted recombination. Only hemizygous males (Y/+,Y/Δ5) and homozygous females (+/+, Δ5/Δ5) were used in these studies and both males and females were included in all experiments. Due to the location of the mutation on the X chromosome, +/+ females and Δ5/Δ5 females were not able to be generated from the same litter but were generated from breeding male Y/+ or Y/Δ5 to female heterozygotes.

B6(C)-H-2Ab1^bm12^/KhEgJ (bm12) and B6.129S2-Cd40lg^tm1lmx^/J (CD40L-KO) were purchased from Jackson Labs and bred in-house. CD40LΔ5 mice were crossed with bm12 mice to create the bm12-CD40LΔ5 line. All animals were caged within the same room. Mice were housed in ventilated micro-isolators under specific pathogen-free conditions in a Rutgers University mouse facility and used at 6–10 weeks of age in accordance with NIH guidelines and under an animal protocol approved by the Animal Care and Use Committee of Rutgers University. Rutgers University is fully accredited by the Association for Assessment and Accreditation of Laboratory Animal Care (AAALAC International) and abides by all applicable laws governing the use of laboratory animals.

### Induction of lupus-like disease by pristane

Mice received a single intraperitoneal injection of 500 μL of the hydrocarbon oil pristane (2,6,10,14-Tetramethylpentadecane; Sigma). Spleen and bone marrow cellular extracts were collected and processed for flow cytometric analysis as previously described (26). In some cases, spleen and kidneys were dissected and immediately embedded and snap frozen with chilled isopentane in Optimal Cutting Temperature compound (Sakura Fintech) and stored at-80 °C.

### Measurement of autoantibody responses

Relative levels of autoreactive antibodies were assayed by indirect ELISA of serum obtained 6 months post-pristane injection or at 2, 4, and 6 weeks after induction of bm12-cGVHD as follows. Nunc Maxisorp 96-well plates were coated with one of four chosen autoantigens in PBS: double-stranded DNA at 5 ug/mL (Thermofisher), nucleosomes at 1 ug/mL, small nuclear ribonucleoprotein complexed with Smith antigen (RNP-Sm) at 1 ug/mL, and ribosomal phosphoprotein 0 (RiboP0) at 1 μg/mL (all three purchased from Surmodics). For dsDNA, plates were first coated with 0.01% poly-L-lysine in PBS. After washing with TBS-T, 50 μL of 1:100 dilutions of serum in PBS + 2% BSA were added and incubated for 2h at 37 °C, followed by washing and addition of AP-conjugated detection antibodies against mouse IgG (Southern Biotech) at 1:1000 in PBS + 2% BSA. Detection was carried out with 4-nitrophenyl phosphate substrate as per manufacturer’s instruction (Sigma S0942) to collect raw OD values at 405 nm.

### Antibodies and Primers

All antibodies and primers used in this study are listed in Supplementary Table 1.

### Immunofluorescent analysis of immune complex deposition in the kidneys

12 μm thick sections of snap-frozen kidneys or spleens from pristane-injected or PBS-injected mice 6-8 months post-injection were prepared in a Leica cryotome. Slides were fixed with ice-cold acetone at-20 °C for 5 min and rehydrated with 1X PBS. Sections were then blocked with 2% BSA in PBS with 1% FcγR block (BioLegend) and 1% rat serum (eBioScience). Kidney samples were stained with anti-mouse IgG–AF647 (1:100) overnight at 4 °C. Excess antibody was washed off with TBS-T. Slides were then counterstained with 2 ug/mL DAPI (MilliporeSigma) and mounted with ProLong Gold Antifade Mountant (Life Technologies). Images for quantification of immunocomplex deposition were taken on an Olympus BX63 Widefield Fluorescent microscope at 20X magnification and analyzed using ImageJ FiJi. 3-5 glomeruli were analyzed per optical field; 3-5 optical fields were captured from both kidneys for 5 mice of each experimental group.

Representative images were taken on a Leica SP8 laser scanning confocal microscope with Leica 10X/NA0.30 HC PL FLUOTAR (lateral resolution: 564) and Leica 40X/NA1.30 HC PL APO oil (lateral resolution: 230) objectives, using a White light laser (460-670 nm) with laser lines 405 nm (DAPI), and 647 nm (AF647) on single z-planes. Leica Application Suite X software was used to linearly adjust brightness and contrast.

### Flow Cytometry

Single cell suspensions (1-5 × 10^6^ cells) from spleen, inguinal and axillary lymph nodes, and bone marrow flow through were stained with Zombie-NIR fixable viability dye (BioLegend) for 15 min, washed and suspended in PBS with 1% BSA, 2% rat serum and 1µg/mL of TruStain FcX anti-CD16/32 (BioLegend) to block FcR binding. After 20 min of incubation on ice, the appropriate primary antibodies conjugated to different fluorescent markers were added at a concentration of 1-10 μg/ml and incubated for 30 min on ice. The cells were washed three times with PBS + 1% BSA then fixed in PBS + 1% paraformaldehyde. When appropriate, fixed cells were permeabilized with 0.2% saponin in PBS + 1% BSA for 10 min and maintained for the duration of cytoplasmic staining. Intranuclear antigens were stained using the True-Nuclear Transcription Factor Buffer (BioLegend) following manufacturer’s protocol. For intracellular cytokine staining, cells were incubated with Cell Activation Cocktail in the presence of 5 μg/mL of Brefeldin A (both from BioLegend) for 4-5 hours before surface staining, fixation, and intracellular staining with saponin. For analyzing DCs, total cells were activated with the Cell Activation Cocktail in the presence of Brefeldin A for 4 hours, followed by surface staining for negative markers (CD3, CD19, CD49b, Gr-1) and positive staining with CD11c and MHC-II. Cells were fixed with 1% PFA for 10 min, permeabilized with saponin as above, and stained with antibodies against IL-6 and IL- 12p40. All data was acquired on a 3-Laser Cytek Aurora cytometer and results analyzed with FlowJo software (TreeStar). FSC-SSC gating for single lymphocytes excluding cell aggregates, small erythrocytes, and dead cell debris, was used for analyzing flow cytometric data. Gates were set using fluorescence-minus-one (FMO) controls.

### Immunoblotting

B cells were isolated from mouse spleens with the MojoSort Mouse Pan B Cell Isolation Kit II (BioLegend) following the manufacturer’s instructions. Cells were processed as previously described (60). Cell equivalents (2×10^6^) cells were separated by SDS-PAGE followed by transfer onto nitrocellulose membranes and processes and described previously. Images were captured and analyzed on a LICOR Odyssey Fc imager. Signals were normalized to B-actin.

### Induction of cGVHD by adoptive transfer of CD4 cells

Spleens and inguinal and axillary lymph nodes were dissected from bm12-WT or bm12-Δ5 mice in sterile conditions and processed into single-cell suspensions. CD4+CD45.2+ cells were subsequently purified using the MojoSort Mouse CD4 T Cell Isolation Kit (BioLegend) following the manufacturer’s protocol and stained with 5μM of CellTrace CFSE Cell Proliferation Kit (Life Technologies) for 20 min at 37 °C in the dark. A lupus-like response driven by chronic graft- versus-host disease (cGVHD) was induced by intraperitoneal injection of 15×10^6^ bm12-WT or bm12-CD40LΔ5 CD4+ cells into corresponding CD40LWT or CD40LΔ5 mice. Control mice were grafted with 15×10^6^ MHC-WT CD4+ cells. Grafting efficiency was verified by flow cytometry on circulating CFSE+CD45.2+ cells collected from submandibular bleeds 3 days post-grafting. Two weeks later, blood and spleens were collected from mice and analyzed for autoreactive antibodies and different lymphoid populations, respectively. To analyze DCs using multiparameter flow cytometry, spleens were collected 5 days post adoptive transfer corresponding to the time when DCs are most active as APCs.

### Dendritic cell and CD4+ cell co-culture

To establish co-cultures of DCs and allogeneic CD4+ T cells, pan DCs were isolated by magnetic sorting from WT and CD40LΔ5 B6 mice and CD4+ T cells were isolated with negative magnetic isolation from bm12-WT and bm12-CD40LΔ5 mice (BioLegend). 1x10^6 DCs were co-cultured with 2x10^6 CD4+ T cells in complete RPMI supplemented with steroid-stripped FBS with or without added 17-β-estradiol (E2) (10 nM; Sigma-Aldrich) for 5 days at 37 °C. Brefeldin A was added in the last 4 hours of culture. Cells were collected and stained for intracellular cytokine analysis by flow cytometry as described above.

### Statistical analysis

Statistical analysis was performed using two-way ANOVA followed by multiple comparisons testing with Tukey’s correction with α = 0.05 only on WT and CD40LΔ5 groups. n.s: p>0.05; *: p<0.05; **: p<0.005. All bar graphs depict arithmetic mean; error bars represent standard deviation. Analysis was done on GraphPad Prism.

## Results

### Increased disease severity in female pristane-injected CD40LΔ5 mice

Experiments were initiated to analyze the effect of modulating CD40L expression through deregulated mRNA stability on PIL mice. For these experiments we analyzed two hallmark disease characteristics previously shown to develop in these mice: 1) the presence of autoreactive antibodies in the serum and 2) antibody-antigen complex deposition in the kidney glomeruli (61). At 24 wks after injection of pristane, antibody titers were measured for anti- Ribosomal phosphoprotein 0 (RiboP0) and anti-RNP complexed with Smith antigen (RNP/Sm), which have been reported to be for the most prevalent autoantibodies produced in C57BL/6-PIL mice (55). As controls we used age-matched, untreated (naïve) B6 mice as well as age-matched and pristane-injected CD40L-KO B6 mice. In control cohorts at the experimental end point, autoreactive antibodies against both antigens were either undetected or at, or slightly above, the experimental threshold for detection. In contrast, CD40LΔ5 male mice showed a strong trend towards decreased RiboP0 antibodies that was highly reproducible but did not reach statistical significance (Fig. 1A, left panel). Also, this response was selective since there was no change in the anti-RNP-Sm response compared to WT males (right panel). In contrast, we found no measurable difference in the RiboP0 response in female cohorts however, the anti-RNP-Sm response in CD40LΔ5 females was unexpectedly elevated over WT females by approximately 30% (Fig. 1A).

**Figure 1:**
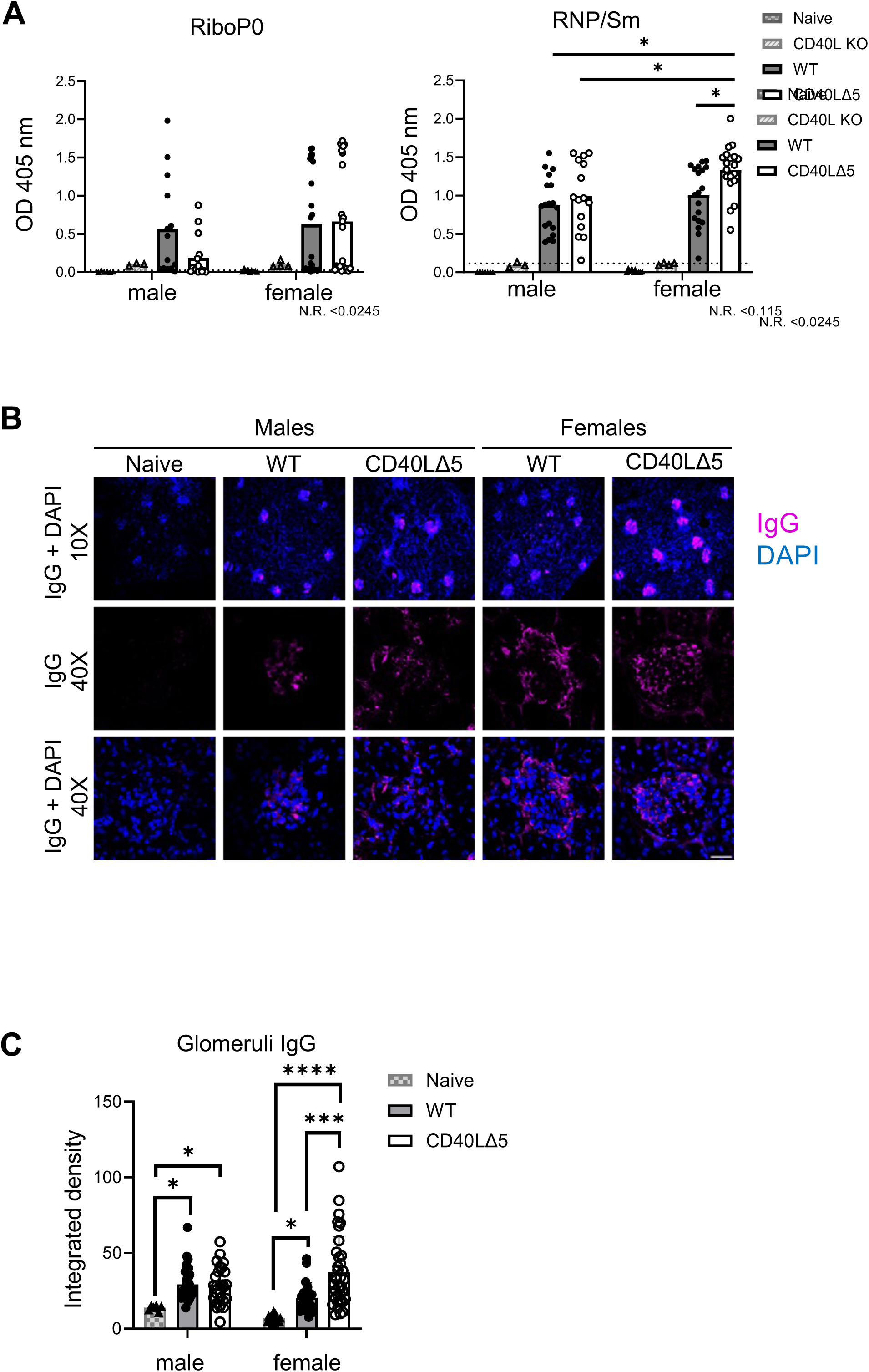
Severity of symptoms in PIL are influenced by levels of CD40L in a sex-dependent manner. (A) Sera from naïve (PBS-injected), and pristane-injected CD40L-deleted (CD40L-KO), WT and CD40LΔ5 mice were measured by indirect ELISA for antibodies against ribosomal phosphoprotein 0 (RiboP0) and ribonucleoprotein complexed with Smith antigen (RNP/Sm). Samples from the WT and CD40LΔ5 groups presenting with a signal below 3X standard deviation (SD) of the naïve groups were labelled as ‘non-responders’ (N.R.; dashed line) and removed from further analysis. (B) Kidney slices from naïve and PIL mice were stained with AF647-conjugated anti-IgG and DAPI, and confocal images captured at 10X and 40X magnification with 2.0 digital zoom. Scale bars at 10X = 150 μm; 40X = 25 m. (C) Widefield images were analyzed with ImageJ to quantify IgG signal per glomerulus. 3-5 glomeruli were analyzed per optical field; 3-5 fields were visualized from both kidneys of 5 mice per group. Histogram represents mean ± SEM integrated density per glomerulus with background signal subtracted. Significance was determined by two-way ANOVA with multiple comparisons using Tukey’s correction: * p≤0.05, ** p≤0.01, *** p≤0.001, **** p≤0.0001

We next asked whether there were changes in kidney pathology that corresponded to the observed CD40LD5-deregulated antibody responses. To this end, kidneys were isolated from both naïve and PIL mice that had been injected 24 wk prior to harvest and stained with anti-IgG antibodies. IgG deposits were observed in male and female PIL WT and CD40LD5 mice but were clearly absent in naïve mice (Fig. 1A). An analysis and quantification of IgG deposits within glomeruli revealed no significant difference in the number of IgG+ glomeruli between WT male, CD40LΔ5 male, and WT female mice although they all showed an increase relative to control naïve mice. However, there was a highly significant increase of approximately 80% in the incidence of IgG deposits in glomeruli from CD40LΔ5 female mice over all other cohorts (Fig. 1B). Taken together, these results show that there is an absolute requirement for CD40L in the development of PIL since CD40L-KO mice exhibited no autoreactive antibodies and importantly, reducing CD40L through a pathway of regulated mRNA stability results in distinctly different outcomes in male and female mice. Whereas a trend towards decreased autoreactive antibodies in male CD40LD5 mice was observed, female CD40LD5 mice displayed a significant increase in antibodies as well as in glomeruli IgG complex deposits; two critical disease indicators of PIL and human SLE.

### PIL- CD40LD5 female mice have an increased pool of splenic ASCs

To extend these observations and determine if corresponding changes in splenic lymphoid populations were present in CD40LD5 mice, we first examined the distribution of CD4 and CD8 T cells and of total B220+CD19+ B cells at 24 wk post injection. Similar to our findings with CD40LD5 mice immunized with either NP-KLH or SRBCs, there were no consistent differences in the T cell compartments between CD40LΔ5 and WT mice (Fig. 2A and Supplementary Fig. 1A) (26). However, analysis of total B cells revealed similar percentages between the two male groups and surprisingly, a lower percentage in the CD40LD5 female mice compared to WT female PIL controls (Fig. 2B and Supplementary Fig. 1B). We also examined the population of CD11c+ aged B cells (ABCs) given their association with heightened disease in multiple models of autoimmunity and found no significant difference in this population (Supplementary Fig. 2).

**Figure 2:**
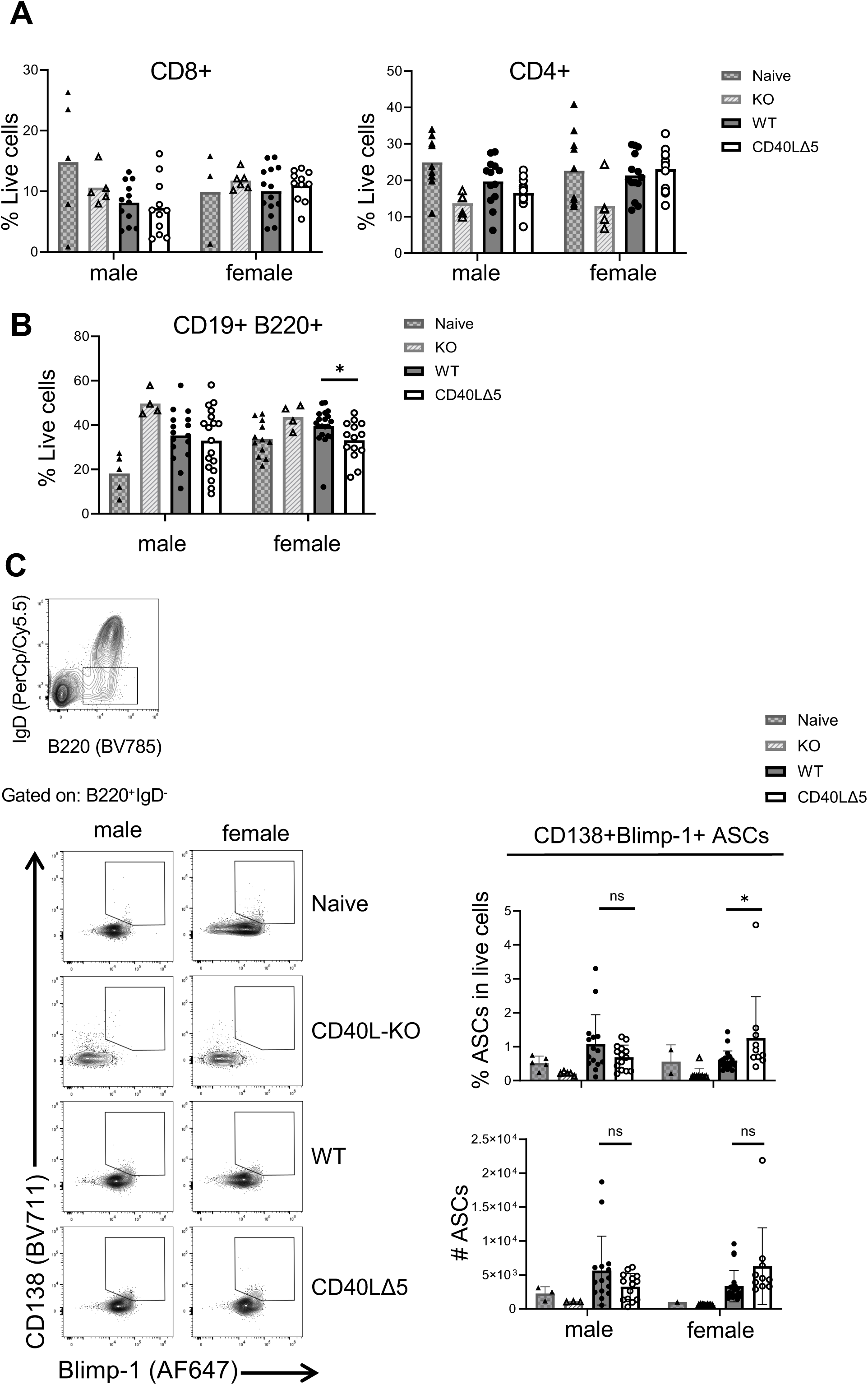
Splenic B cell populations are differentially affected by changes in CD40L expression on CD4 T cells. (A) Compilation of data showing the proportion of CD4 and CD8 T cells and (B) CD19+B220+ B cells in naïve, KO, WT and CD40LΔ5 PIL mice as determined by flow cytometry. (C) Representative analysis of splenic ASCs (IgD-B220+CD138+ Blimp-1+) (left) and associated relative frequencies and total cell counts (right). Data are shown as mean ± SEM from five independent experiments (n=6-14). Significance was determined by two-way ANOVA with multiple comparisons using Tukey’s correction where * p≤0.05.

Assessment of the CD19+IgD-CD138+Blimp1+ antibody producing cells (ASCs) revealed that, as predicted, naïve and pristane-injected CD40L-/- mice had low to undetectable levels. In male CD40LΔ5 mice there appeared to be a strong trend towards decreased numbers and percentages of ASCs compared to male WT mice that did not reach statistical significance but agreed with our antibody data. We also observed a higher proportion of ASCs in the female CD40LD5 mice, a finding highly consistent with the elevated levels of circulating autoreactive antibodies associated with this cohort (Figure 2C).

### The CD40LΔ5 mutation results in elevated numbers of GC B cells in female PIL mice

To further understand the relationship between lowered CD40L expression and the atypical and exacerbated autoimmune response in female mice, we next asked whether there were corresponding changes in the GC response that could be linked to lowered CD40L. Surprisingly, spleens from 24 wk post injection PIL mice showed clear histological evidence of intact GCs in all cohorts with the presence of GC B cells (green) within the CD169+ follicles (Supplementary Fig. 3). This result was clearly different than what was observed in CD40LD5 mice following antigen challenge which resulted in GCs being highly disorganized compared to WT mice (26).

We next evaluated the GC populations by first analyzing the Tfh (CD4+CD8-PD-1+CXCR5+) population and found no significant difference in the percentage of Tfh cells between the different cohorts, except for the CD40L-KO mice for which this population was not detected (Fig. 3A). In contrast, there was a 2-fold enrichment of GL7+ B cells (B220+CD19+GL7+Fas+) in the female CD40LΔ5 PIL mice compared to WT females that was not seen when comparing the male cohorts (Fig. 3B). To determine whether differential CD40 signaling in B cells was the basis for the gender-specific responses we analyzed signaling pathways known to be triggered by CD40 engagement in B cells (1). Immunoblotting was carried out with CD19+ B cells purified from splenic extracts by negative selection and analyzed for activation of p52, p65 and Akt. We found that activation was decreased in all pathways indicating that CD40 signaling is clearly compromised in CD40LD5 mice. However, decreased CD40-mediated signaling was observed in both male and female CD40LD5 B cells suggesting that alterations in activation did not explain the gender-based disparities observed in the PIL disease severity (Fig. 3C).

**Figure 3:**
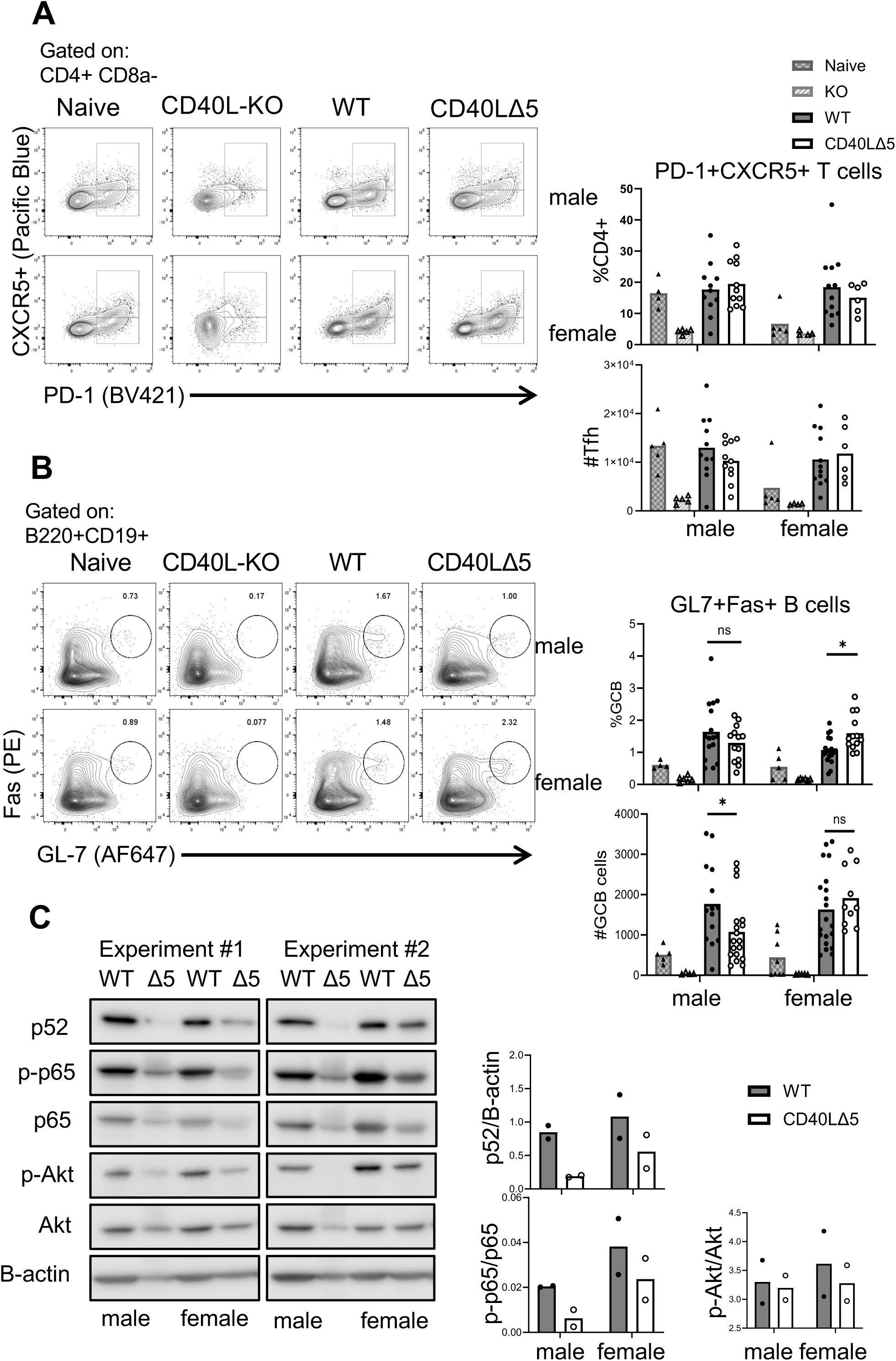
Reduced CD40L results in an increase in GC but not extrafollicular B cells in female mice. (A) Representative analysis of naïve and PIL male and female CD40L-KO, WT and CD40LΔ5 mice showing the identification of splenic Tfh cells (CD4+CD8-PD-1+CXCR5+) 24 wks post pristane injection. Graphical representation of 5 or more independent experiments. (B) Flow analysis of splenic cells to identify B220+CD19+Fas+GL-7+ GC B cells showing naïve and PIL male and female CD40L-KO, WT and CD40LΔ5 mice (left). Relative frequency and total number of GL7+ B cells compiled from greater than 5 independent experiments (5–12). All data are presented as the mean ± SEM with significance being determined by two-way ANOVA with multiple comparisons using Tukey’s correction where * p≤0.05. (C) Data from two independent experiments depicts splenic CD19+ B cells that were negatively isolated from 6-month PIL mice and analyzed by immunoblot for activation of p52, p65 and Akt pathways. Densitometry analysis is depicted on the right and was quantified as follows: for p52, values were normalized to those of B-actin; for phosphorylated p65 and Akt, both were normalized to total levels of protein on the same blot.

Symptoms in PIL mice are evident only after six months and therefore this timing makes it difficult to assess the initiating cellular and molecular events that lead to disease. Because we wanted to track the initiating events that lead to differential autoimmune responses driven by reduced CD40L, we chose to use a second model of lupus that allows for assessing initiating events early in the development of disease.

### Using the bm12-cGVHD model to analyze initiating events in lupus

Bm12 mice harbor a three nucleotide/three amino acid change in the MHC class II antigen, H-2Ab1^bm12^ and introduction of bm12-CD4+ T cells into MHCII-WT animals on the B6 background induce a chronic graft-versus-host disease that presents with the expansion of autoreactive B cells and leads to a lupus-like disease driven by autoreactive antibodies (49, 56, 62). This model is distinct from the PIL model since changes in immune cell profiles are evident between 7- and 14-days following cell transfer (49, 59, 62–64). Furthermore, disease is primarily mediated by CD4+ T cell-B cell interactions, particularly between Tfh-GL7+ B cells, as both populations observe significant expansions shortly after grafting (49, 56, 62).

To prevent complications arising from a mixed population of WT and CD40LD5 CD4+ T cells within the same mouse, we matched both gender and CD40L genotypes in donor and recipient mice creating CD40LD5-bm12 mice. Specifically, under these conditions, CD40L deficiency would be present during host B cell ontogeny as well as following allogenic activation with the transfer of bm12-CD4+ T cells. Additionally, to enable monitoring of the behavior of the grafted CD4+ T cells, we initially used the congenic CD45.1 and CD45.2 alleles to identify the behavior of transferred cells. To this end, we crossed our CD45.1/CD40LΔ5 line with CD45.2/H-2Ab1^bm12^ mice on the C57BL/6 background to generate a ‘bm12Δ5’ mouse carrying the bm12 and CD40LΔ5 mutations with the CD45.2 allele (Fig. 4A) (56). Thus, all donor cells from all genotypes carried the CD45.2 marker.

**Figure 4:**
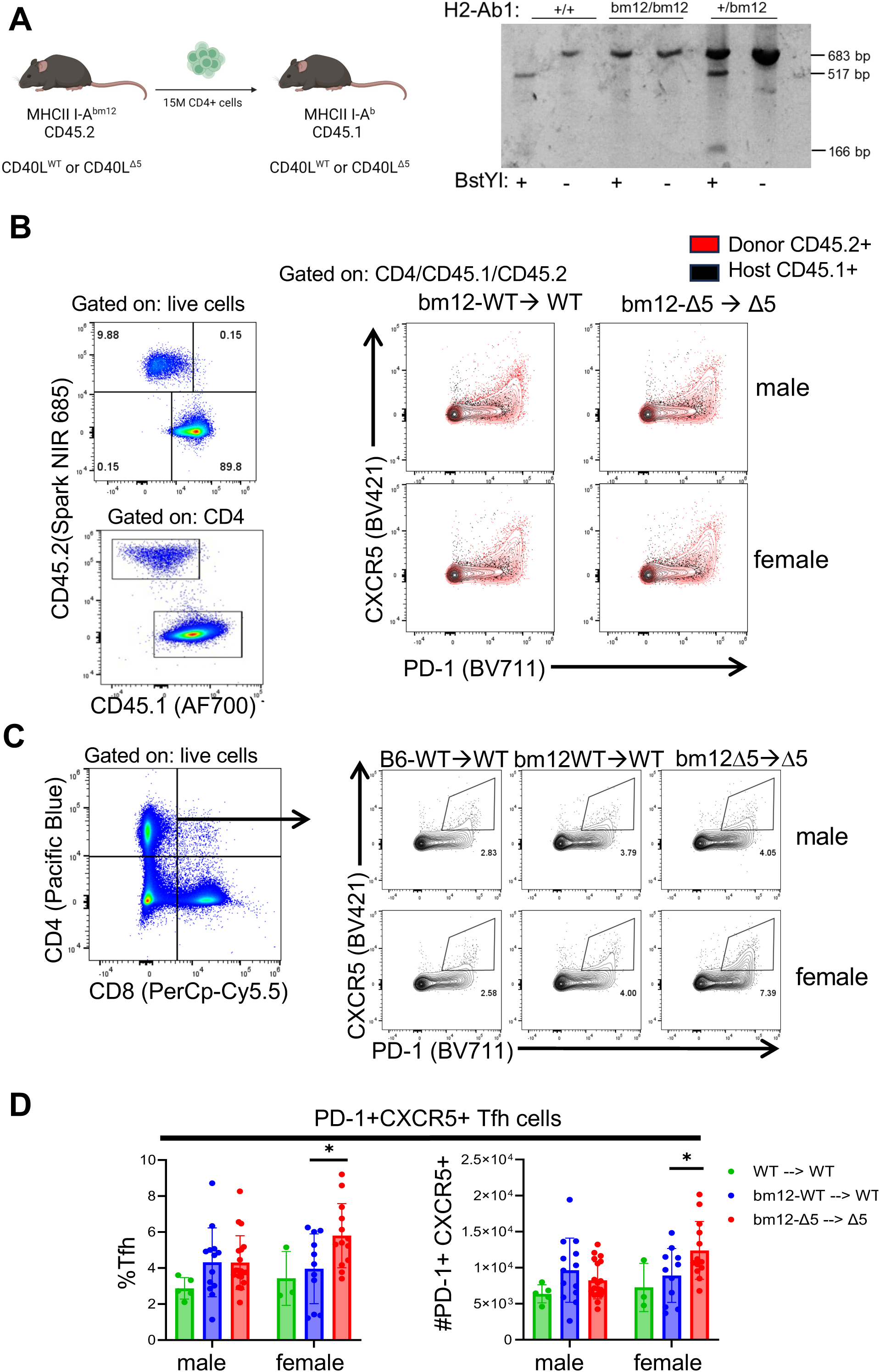
The bm12 model of cGVHD to study the early onset of disease. (A) Graphical summary of disease induction by transfer of allogenic CD4+ cells (left) and representative PCR genotype showing the amplified bm12 region of the H2-Ab1 gene from genomic DNA. PCR samples were digested with BstYI and analyzed by gel electrophoresis (right). (B) (left) Representative flow analysis confirming the presence of donor bm12 CD4 T cells into matched recipients. (Right) Representative histograms of CD4+CD45.1+CD45.2+ cells analyzed for the presence of Tfh cells (PD-1+CXCR5+) taken from host mice 2 weeks after transfer; host CD45.1+ (black) and donor CD45.2+ (red) CD4+ populations are shown. (C) A representative analysis showing identification of male and female Tfh cells using PD-1 and CXCR5 following transfer of bm12WT (middle histograms) or bm12Δ5 CD4 T cells (right histograms) into WT or CD40LΔ5 gender-matched recipients. Control transfers of male and female WT CD4 T cells into sex-matched WT recipients are shown in left histograms. (D) Compilation of data from at least five different independent experiments (n=5-14) shown as the mean ± SEM. Significance was determined by two-way ANOVA with multiple comparisons using Tukey’s correction: *p≤0.05.

Two weeks after adoptive transfer of bm12 CD45.2+ CD4 T cells into male and female WT and CD40LD5 recipients we observed that PD-1 and CXCR5 expression was upregulated on a marked proportion of CD45.2+ cells indicating that transferred cells had acquired a Tfh phenotype (Fig. 4B, red dots). Because of evidence that recipient Tfh cells are important for “nurturing” of B cells during development of disease and participate in the bm12-induced response, all Tfh cells were analyzed in subsequent experiments (56). Importantly, we saw an enrichment of Tfh cells in all cohorts receiving bm12 CD4+ T cells over those receiving WT donor cells indicating that the Tfh population was expanding due to MHC incompatibility (Fig. 4C).

When all experimental groups were compared against each other, there was a significantly higher percentage of Tfh cells in host female CD40LD5 mice receiving bm12-CD40LΔ5 CD4 T cells (bm12Δ5 ◊Δ5) compared to all other groups (Fig. 4D).

### Enhanced autoimmune responses are detected in CD40LΔ5 female mice

To further evaluate the bm12-induced immune response we analyzed antibodies against four different antigens identified as the primary autoantibodies produced in the bm12-cGVHD model (56, 57). Consistent with our results with PIL mice, a significant increase in all groups of autoantibodies was detected in recipient CD40LΔ5 females (bm12Δ5 ◊Δ5) compared to WT female mice (bm12WT◊WT) with no observed differences between the male cohorts (Fig. 5A). This striking difference in autoantibody production was also reflected in the increased proportion of CD19+IgD-CD138+Blimp1+ ASCs in B cells from CD40LΔ5 females (bm12Δ5◊Δ5) compared to WT females (Fig. 5B).

**Figure 5.**
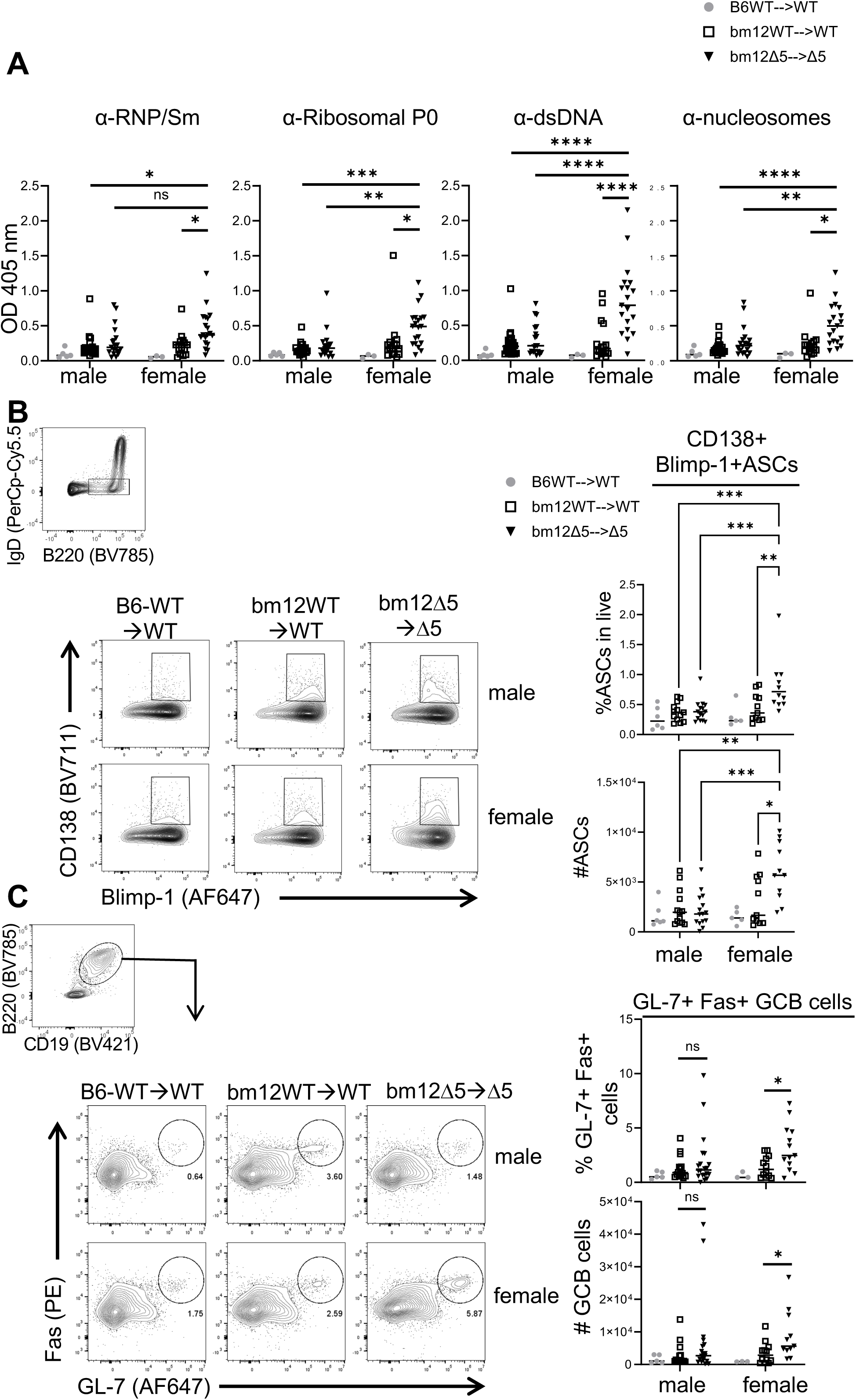
Autoantibodies and antibody-secreting cells are significantly elevated in female mice expressing CD40LΔ5 mutation following bm12-cGVHD induction. (A) Levels of IgG autoantibodies in serum against RNP/Sm, and Ribo P0, dsDNA, and nucleosomes were quantified by indirect ELISA at 2 weeks post transfer of bm12-CD4+ T cells. (B) Representative FACS analysis of splenic B220+IgD-CD138+ Blimp-1+ ASCs (left) and compiled data in right graphs. (C) B220+ CD19+GL7+Fas+ splenic GC B cells and representative histograms for recipient male and female WT mice receiving WT (left) or bm12WT (middle) CD4 T cells and CD40LΔ5 recipient mice receiving bm12Δ5 CD4 T cells (right). Quantification of relative frequencies and total cell counts are shown on the right. Data reflects the mean ± SEM from at least five independent experiments with 3-12 animals per experiment. Significance was determined by two-way ANOVA with multiple comparisons using Tukey’s correction: * p≤0.05, ** p≤0.01, *** p≤0.001, **** p≤0.0001.

Notably, the bm12-cGVHD model is highly dependent on GC activity and the associated B cell development mediated by donor bm12 CD4+ T cells. Specifically, adoptive transfer of bm12 CD4+ T cells into non-bm12 mice leads to the activation of the grafted cells, the induced expression of T follicular markers, PD-1 and CXCR5, and the subsequent migration of CD4+ T cells into the follicles of secondary lymphoid organs (56, 65). To understand the impact of the CD40LD5 mutation on GC development in this model, we analyzed GL7+ B cells from recipient mice two weeks post transplantation. We observed a significant enrichment of GL7+ B cells in bm12-grafted CD40LΔ5 female mice compared to WT female mice. Also, there were no significant difference either between the recipient female WT mice and the two recipient male cohorts (Fig. 5C). Together, this data shows that the recipient CD40LΔ5 females displayed a much more robust autoantibody response as well as enhanced numbers of Tfh cells, ASCs and GL-7+ GC B cells suggesting that the gender disparity associated with reduced CD40L expression is not defined by a specific model of lupus and that this disparity manifests as an early stage in the development of disease.

### CD4+ inflammatory cytokines are highly prevalent in bm12-cGVHD female Δ5 mice

Although we were unable to observe changes in CD40 signaling that possibly underlie the gender-and CD40LD5-specific responses in PIL mice, we wanted to re-evaluate these signaling pathways to determine if there were effects on these pathways in recipient mice at an earlier time point in disease progression. However, we were again unable to observe changes in B cell signaling pathways that corresponded to gender-specific responses (Supplementary Fig. 4). Therefore, we chose to focus on other extrinsic signals that could affect the unique responses observed in the CD40LD5 female mice. Given that the cytokine milieu plays a critical role in the development and amplification of disease in both lupus models and human SLE patients, our initial focus was on evaluating CD4+ T cells for cytokines implicated in disease initiation and progression (66–69). Following isolation and adoptive transfer of bm12 CD4+ T cells into recipients, splenic cells were analyzed at 2 weeks post transfer for intracellular expression of cytokines IFN-γ, IL-2, IL-4 and TNF-α. In comparing male mice, we found no significant difference in the percentage and number of cytokine-producing CD4+ T cells. In contrast, we observed a significant decrease in the percentage of TNF-α expressing CD4+ T cells as well an increase in the percentage of CD4+IL-4+ cells in CD40LΔ5 female mice relative to WT females (Fig. 6A). Also, determination of Th1 (CD4+ IFN-γ+ IL-2+) and Th2 (CD4+ IFN-γ- IL-4+) subsets revealed that WT and CD40LD5 males and WT females were enriched in Th1 cells compared to CD40LD5 females. In contrast, there was a higher proportion of Th2 cells in CD40LD5 females relative to the other three groups and this represented a decreased Th1/Th2 ratio (Fig. 6B). Importantly, these results suggest that the cytokine milieu produced by the donor CD4+ T cells may be essential for driving the gender-specific and CD40L-dependent disparities observed in the bm12-cGVHD model.

**Figure 6.**
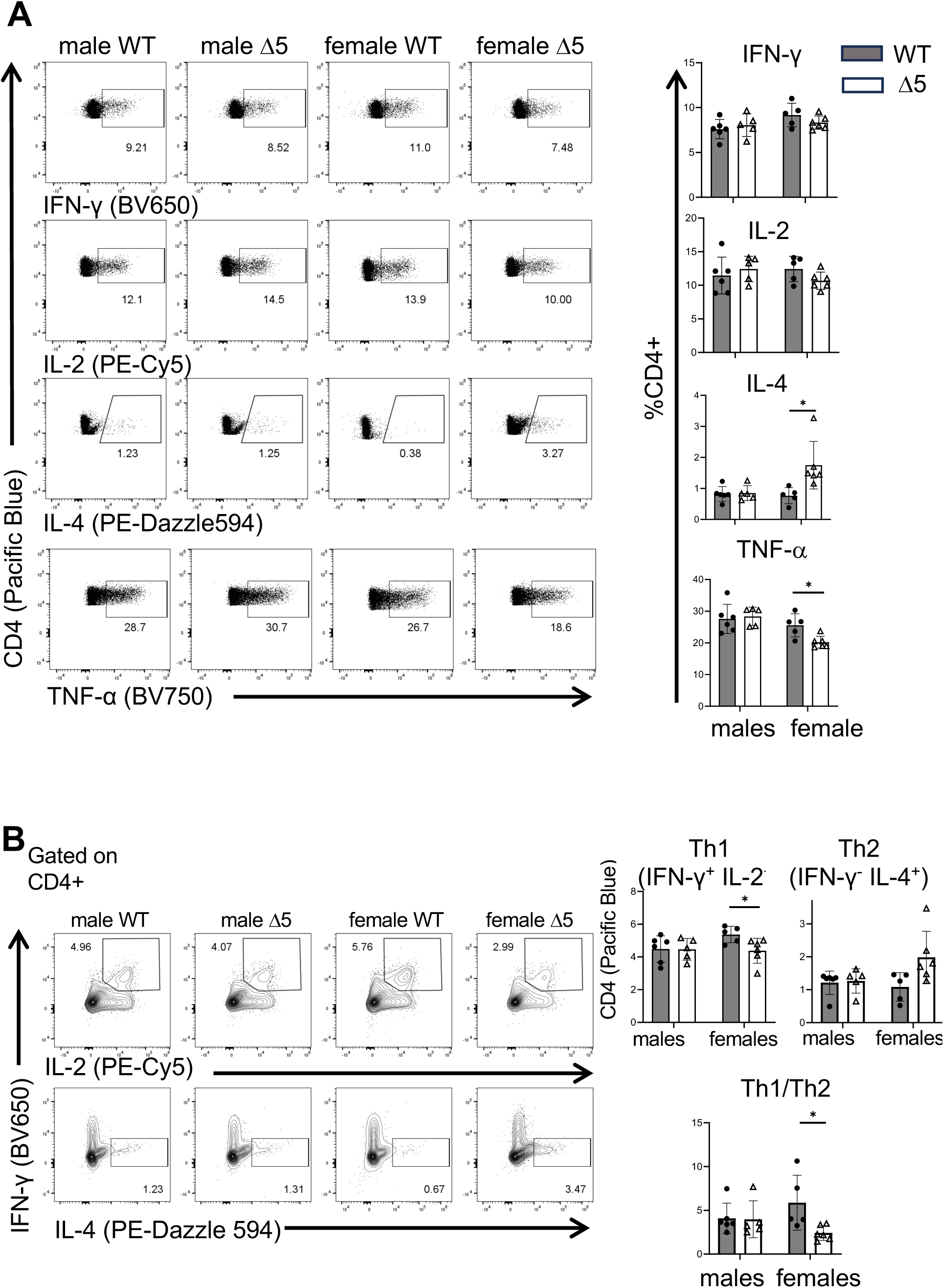
Reducing CD40L expression in female mice changes the cytokine profile in CD4+ T cells. (A) Representative histograms showing splenic CD4+ T cells from male and female WT and CD40LΔ5 mice that had received bm12-CD4 WT or bm12-CD40LΔ5 CD4 T cells 2 weeks earlier. Splenic cells were isolated and analyzed by intracellular staining and flow cytometry for the expression of IFN-γ, IL-2, IL-4, and TNF-α. Graphical representation of two or more independent experiments showing the frequency of cytokine positive CD4+ T cells following activation with PMA and ionomycin for 4-5 h in the presence of brefeldin A (right graphs). (B) Analysis of male and female populations of Th1 (IFN-γ+ IL-2+) and Th2 (IFN-γ-IL-4+) cells in control (B6WT->B6WT), bm12-WT->WT and bm12-Δ5-CD40LΔ5 mice. Representative histograms are shown on the left and on the right graphs of compiled data from three independent experiments showing the frequency of cytokine-expressing CD4+ T cells from the different cohorts. The ratio of the percentage of Th1/Th2 cells are shown in the bottom graph. Statistical significance was determined by two-way ANOVA with multiple comparisons using Tukey’s correction: * p≤0.05.

### Expression of IL-12 is decreased in DCs from female CD40LΔ5 mice

Because DCs play such a central role in establishing the cytokine milieu following engagement with activated CD4+ T cells, we sought to examine the DCs in recipient mice to determine if changes in CD40L-specific responses could explain the observed gender-based outcomes in disease parameters. Therefore, splenic DCs were analyzed from recipient mice five days post transfer of purified CD4+ T cells and identified as lineage negative, and MHCII and CD11C positive (CD3^-^ CD19^-^ CD49b^-^Gr-1^-^MHCII^+^CD11C^+^). We found that WT males had a significantly higher DC population relative to all other cohorts suggesting intrinsic differences in WT males and females in the DC population size following initiation of bm12-cGVHD lupus. Also, the CD40LD5 mutation corresponded to a decrease in percentage DCs in males but not in females (Fig. 7A). DCs were further analyzed for expression of intracellular IL-6 and IL-12, two cytokines shown to enhance and limit the development of pathogenic Th2 cells in SLE, respectively (70–72). We examined cells both prior to and after ex vivo activation with PMA and ionomycin and found no significant differences in the percentage or number of IL-6+ DCs between the different cohorts (Fig. 7B). In contrast, analysis of IL-12-expressing DCs revealed that WT female recipients had higher percentages and numbers compared to all other cohorts and the difference was clearly evident with and without ex vivo stimulation. In striking contrast, the CD40LD5 mutation resulted in female mice with a reduced population of IL-12+ DCs compared to wildtype females. (Fig. 7C). Therefore, the CD40LD5 mutation corresponded to a significant decrease in the proportion of IL-12-expressing DCs in female but not male mice.

**Figure 7.**
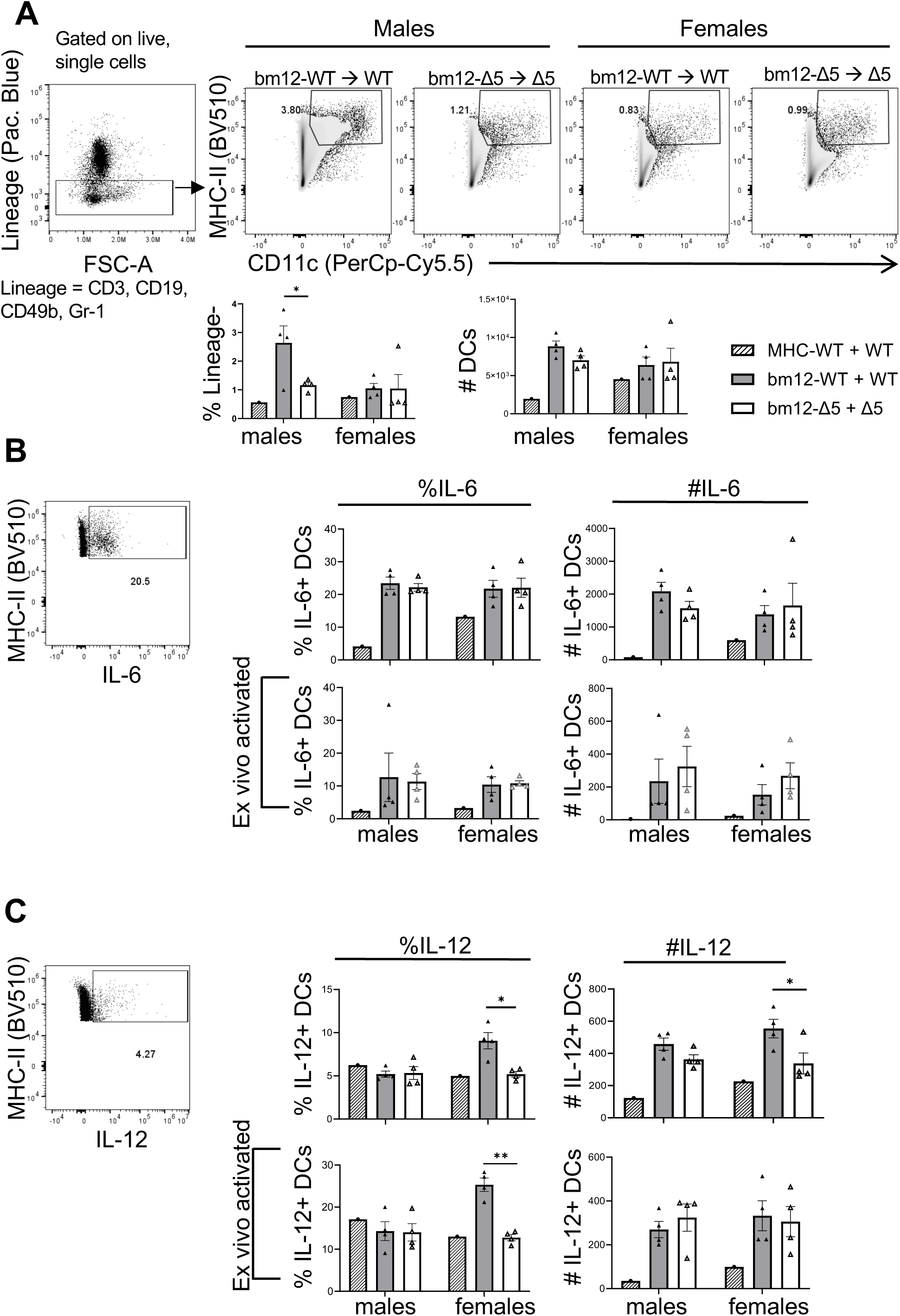
Reduced CD40L results in a differential pattern of cytokine expression only in splenic DCs from female mice. (A) Representative gating of lineage (Lin) negative (CD3-, CD19-, CD49b-, F4/80-), CD11c+ MHC-II+ splenic DCs isolated from male and female B6 or B6-CD40LΔ5 mice receiving either bm12-WT or bm12-Δ5 CD4 T cells 5 days prior to sacrifice. DCs, presented as the percentage of Lin-cells, are shown in the graphs below. (B) Intracellular cytokine detection by flow cytometry in Lin-CD11c+ MHC-II+ DCs. Gating of positive cells is shown on the left and on the right the quantification for cells taken directly from the mice (top panels) and cells cultured *ex vivo* with PMA and ionomycin for 5h in the presence of Brefeldin A. Significance was determined by two-way ANOVA with multiple comparisons using Tukey’s correction: * p≤0.05, ** p≤0.01

### Exogenous estrogen does not change the cytokine profile of male and female CD40LD5 DCs

Having linked the dimorphic phenotype of the CD40LΔ5 mutation to DC cytokine expression we next asked whether 17-β-estradiol (E2) was a factor in skewing female CD40LΔ5 CD4 T cells toward a Th2 phenotype in the presence of lowered CD40L signaling. In this scenario, we would expect to see a similar pattern of cytokine expression in male CD40LΔ5 DCs stimulated with male CD40LΔ5 CD4 T cells in the presence of E2 leading to a predominant Th2 response. We therefore isolated CD4 T cells from bm12-WT and bm12-Δ5 mice and co-cultured them with isolated DCs from gender-matched WT-B6 or CD40LΔ5-B6 mice in steroid-free media with or without added E2 (73). After 5 days, the expression of DC-specific IL-6 and IL-12 was determined by intracellular staining (Fig. 8A). In cultures containing *male* WT-bm12 CD4 T cells, we found that the number and percentage of IL-6-expressing WT DCs increased approximately 2-fold following E2 treatment. In contrast, there was no observed increase in IL-6-expressing DCs from cultures containing male CD40LΔ5 -bm12 T cells (Fig. 8B (left panel), compare M-E2- and M- E2+). Analysis of co-cultures containing female WT or CD40LΔ5 cells found that the percentage of IL-6-expressing DCs was increased in CD40LΔ5 cultures only and was consistent with the enhanced Th2 response in these cells (Fig. 8B (left panel, compare F-E2- and F-E2+). A similar assessment of IL-12 expression revealed an increase in the proportion of male WT and CD40LΔ5 DCs in the co-cultures following E2 exposure. Notably, the population of female IL-12+ DCs was relatively unchanged following E2 exposure although the proportion in the CD40LΔ5 DC was consistently lower than in WT cultures (Fig. 8C). Therefore, even though there were clear changes in cytokine responses in male CD40LΔ5 DCs with E2 signaling, these signals did not produce a cytokine expression pattern that was similar to female CD40LΔ5 DCs. Importantly, with or without added E2, male and female DCs receiving activation signals via CD40LΔ5 T cells express a cytokine profile that selectively skews CD4 T cells towards a predominant Th1 or Th2 phenotype based on gender and which corresponds to reduced or heightened levels of autoantibody, respectively.

**Figure 8.**
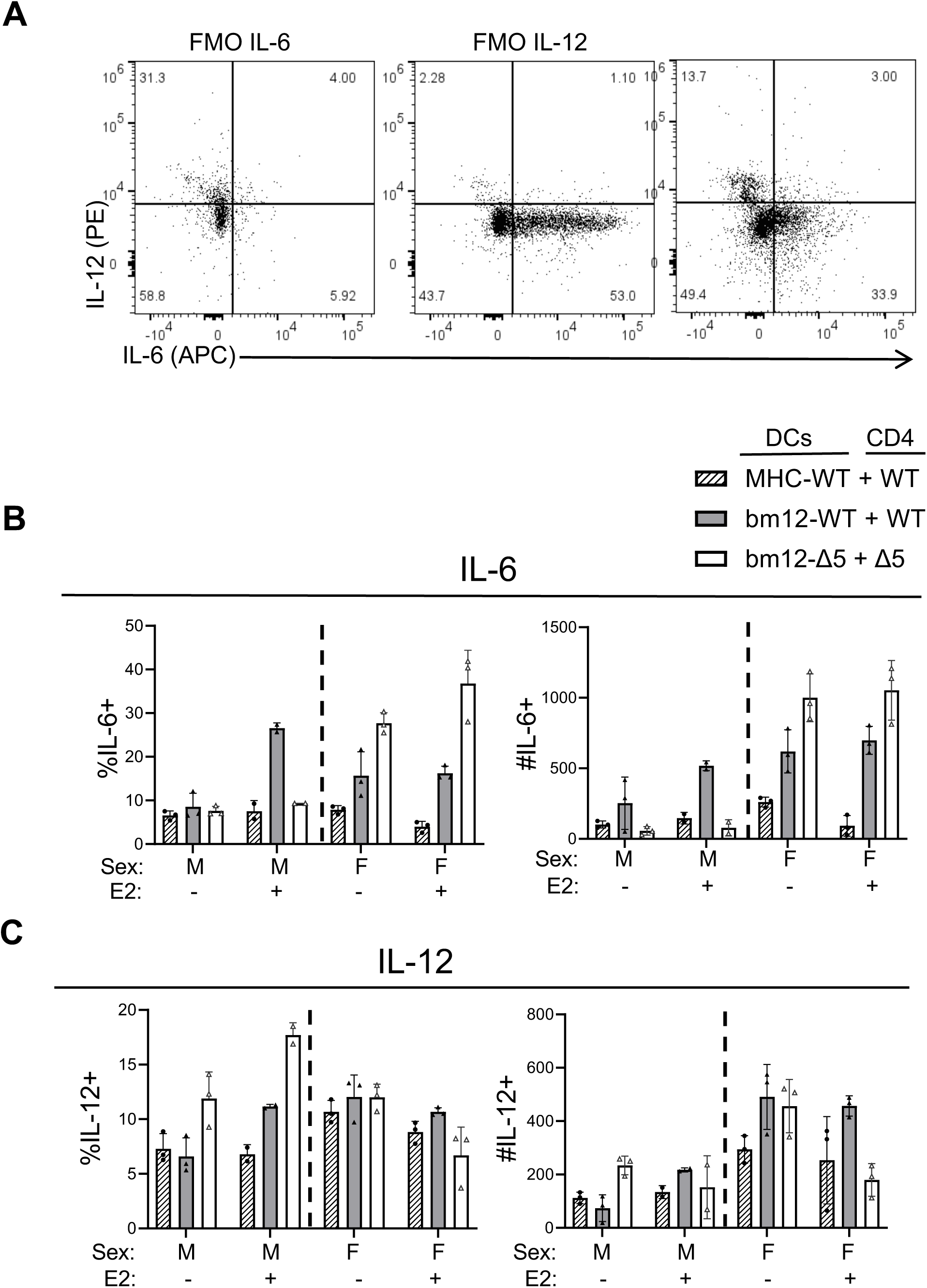
CD40L- and gender-specific expression of cytokines in histoincompatible cultured DCs is independent of estrogen. Splenic DCs from MHCII-WT mice (either B6 or B6-CD40LΔ5) mice were isolated 5 days after adoptive transfer and co-cultured with CD4+ T cells from MHCII-bm12-WT or bm12-CD40LΔ5 mice at a 1:2 ratio in steroid-free media supplemented with 17-β-estradiol (E2). Shown in (A) are representative histograms of IL-6 and IL-12 positive DCs showing control FMO profiles for IL-6 and IL-12 and an E2-stimulated WT culture expressing IL-6 and IL-12 (right). (B and C) Compiled intracellular expression data of (B) IL-6 positive and (C) IL-12p40 positive DC cells showing both the frequency (left) and number (right) of positive cells.

## Discussion

In this work we sought to extend our understanding of CD40 signaling in autoimmunity by using mouse models of lupus that express reduced levels of CD40L through a change in mRNA stabilization. Unexpectantly, we uncovered a novel role for optimal CD40L expression that is linked to limiting, rather than augmenting disease in female B6 mice. The finding that male and female mice harboring the same reduced CD40L phenotype have significantly different autoimmune responses was surprising given the abundance of data supporting the role of CD40L as a driver of autoimmunity (40, 50). However, our results were not completely incompatible with this premise since we observed that CD40L-KO mice of both sexes were protected from disease symptoms and more importantly, it was only upon reducing and not eliminating CD40L expression that we observed our novel findings in female mice. Although autoantibodies in female B6-CD40LΔ5 mice were persistently elevated, in male B6-CD40LΔ5 mice they were not only significantly lower than mutant females they also trended lower relative to WT males. This was seen most clearly in the PIL model suggesting that consistent with the predicted function of CD40L in SLE, lowering CD40L-CD40 signaling in PIL male mice resulted in an overall reduction in disease symptoms.

The dimorphic disease indicators associated with the CD40LΔ5 mutation in female mice included increased autoantibodies and ASCs in both models as well as significant increases in Tfh cells, GC B cells and a decreased Th1/Th2 ratio in the bm12-cGVHD model. This profile is consistent with lowered CD40 signaling between CD4 T cells and DCs resulting in reduced IL-12 expression with an associated decrease in the IL-12Rβ- and Tbet-dependent transcriptional program critical for the development of a Th1 response (2, 19, 20, 74–79). Tfh cells induce GC responses that lead to autoantibody production, whereas Th1 and Th17 cells contribute to immunoglobulin switching and lupus nephritis (80–83). The observed increase in Tfh cells is likely connected to the expanded population of ASCs which drives the increase in autoreactive antibodies observed in female B6-CD40LΔ5 mice. In support of our results, increased Tfh cell numbers have been found in patients suffering from SLE and in mice with a targeted deletion of the Ets1 gene in CD4 T cells (*Ets1*^DCD4^). *Ets1*^DCD4^ mice develop an SLE-like disease with a corresponding expansion of Tfh cells and a dominant Th2 response suggesting that suppressing Th2 differentiation limits the onset of SLE (30, 84, 85).

Reduced CD40L in both mouse models gave similar increases in autoreactive antibodies in female mice however the bm12 model also allowed us to analyze very early events attributable to reduced CD40L-CD40 signaling. Specifically, we were able to show that decreased CD40L on allogeneic CD4 T cells reduced expression of IL-12 from female DCs only. Previous reports have shown that within an environment of reduced or absent IL-12, CD4 T cells differentiate preferentially into Th2 cells establishing a more robust disease phenotype at early steps in its initiation (35, 86). Also, CD4+ T cells default to a Th2 pathway in the absence of positive signals (such as IL-12) that drive differentiation into other Th cell lineages (87). The CD40LΔ5 model reduces CD40L expression to approximately 60% of WT levels. However, it remains unclear why male and female DCs display such different cellular responses to a similar degree of reduced expression.

One interesting observation that supports the idea that inherent sex differences exist in DCs at an early point of engagement, is that at 5 days following transfer of alloreactive T cells, differences in the levels of DC-induced cytokines were clearly evident in WT B6 male and female mice receiving WT-bm12 T cells. Notably, the overall DC population was increased in male WT mice relative to all other cohorts whereas the IL-12-expressing DC population was specifically expanded in female WT mice. Given that we have been unable to detect any differences in the expression pattern of CD40 on male and female DCs suggests that even in WT recipients, similar levels of CD40 signaling lead to gender-specific changes in IL-12 expression. Thus, DCs appear to be differentially programmed for activation responses based on inherent sex differences prior to or concurrent with receiving a CD40 signal via allogeneic CD4 T cells.

Our focus on estrogen as a possible candidate for causing differential responses in DCs was based on numerous studies showing the role of estrogen in immune and autoimmune responses in both mice and humans (88). Estrogen is required for both the differentiation of DCs from bone marrow progenitor cells and for the optimal activation and DC expression of MHCII, CD86 and CD40 (89–93). Also, the contribution of hormonal status to DC homeostasis is underscored by the fact that multiple hormones, including estrogen and progesterone, can modulate immune cells and DC differentiation, maturation and function (94). Both male and female DCs express E2alpha (E2a) and E2beta(E2b) receptors however, previous work has been conflicting as to whether there is both a direct connection between estrogen and the production of IL-12 in CD11C+ cells (95, 96). Although manipulation of estrogen levels has been shown to alter DC function, specific different levels between males and females during autoimmunity have not been reported. In contrast, some infection models have indirectly pointed towards sex-based differences in DC function due to dimorphic expression of immune response genes or other proteins normally expressed by DCs (97). Importantly, we found that exposure to E2 in vitro, did not convert male responses to female responses, again suggesting that sex-based intrinsic differences in DCs exist prior to or concurrently with bm12-WT or-CD40LΔ5 T cell contact and CD40L-CD40 signaling.

Our working model for understanding the basis for observed sex-specific differences in our models is that there is a threshold of CD40 signaling driven by allogenic or autoreactive CD4 T cells required to instruct DC-specific cytokine expression and this signaling threshold is notably different for female and male DCs. The cytokine milieu dictates the differentiation of CD4 T cells into effector subsets. At the very early stages of DC and bm12-CD4 engagement, a Th1 response is more protective and a predominant Th2 response more favorable for disease development.

Importantly, the roles of cytokines can differ during the course of disease. For example, IL-12 has been shown to function in a positive feedback loop with IFNγ, promoting plasma cell differentiation and suppressing GC formation (98). The difference in threshold response is similar to what we observed with the CD40LΔ5 model in response to a TD immune response (NP-KLH).

In this case, different CD40-dependent processes, such as B cell memory formation, were highly sensitive to the drop in CD40 signaling levels whereas other processes (i.e. class switching and development of plasmablasts) were less so. Notably we did not detect sex-specific responses modulated by decreased CD40L in these immune response models (26).

In conclusion, our unexpected and novel findings suggest that optimal CD40L expression plays a fundamental role in modulating early cellular responses to initiating factors that lead to SLE. Importantly, they support a nuanced relationship between the level of CD40L expression and the progression of disease that is sex dependent. Defining how CD40 signaling is integrated into the different cellular responses of this complex and devastating disease will increase our overall understanding of gender as a major disease risk factor and support the incorporation of this knowledge into new and future treatments.

We thank Dr. Ping Xie for critically reading of this manuscript and for their many helpful discussions and insight. We acknowledge the many past and present lab members of the Covey lab for valuable contributions to this project. This work was supported by an AAI training grant to DPM and by grants from the National Institutes of Health (AI-57596 and AI-107811) to LRC.

