## Supplemental Figs for "Lowering CD40L expression in Murine Lupus Results in an Increase in Disease Indicators in Female but not Male B6 mice"

**Supplementary Table 1: List of antibodies**

| **Table 1: Antibodies** | | | | |
|---|---|---|---|---|
| **Antigen** | **Fluorophore** | **Clone** | **Source** | **Catalog #** |
| *T cell panel* | | | | |
| CD4 | Pacific Blue | GK1.5 | BioLegend | 100428 |
| CD8 | PerCp-Cy5.5 | 53-6.7 | BioLegend | 100734 |
| PD-1 | BV711 | 29F.1A12 | BioLegend | 135231 |
| CXCR5 | BV421 | L138D7 | BioLegend | 145511 |
| CD69 | AF700 | H1.2F3 | BioLegend | 104539 |
| CD25 | BV510 | PC61 | BioLegend | 102041 |
| CD40L | APC | MR1 | BioLegend | 106510 |
| FoxP3 | PE | MF-14 | BioLegend | 126404 |
| IL-10 | PE-Dazzle 594 | JES5-16E3 | BioLegend | 505033 |
| CD40L | PE | MR1 | BioLegend | 106505 |
| CD45.2 | Spark NIR 685 | 104 | BioLegend | 109863 |
| CD45.1 *or* | AF700 | A20 | BioLegend | 110723 |
| *GC B cell panel* | | | | |
| CD19 | BV421 | 6D5 | BioLegend | 115538 |
| B220 | BV785 | RA3-6B2 | BioLegend | 103246 |
| Fas | PE | SA367H8 | BioLegend | 152607 |
| GL-7 | AF647 | GL7 | BioLegend | 144606 |
| IgM | PE-Dazzle 594 | RMM-1 | BioLegend | 406530 |
| IgD | PerCp-Cy5.5 | 11-26c.2a | BioLegend | 405710 |
| IgG1 | BV650 | RMG1-1 | BioLegend | 406629 |
| IgG2b | PE/Cy7 | RMG2b-1 | BioLegend | 406714 |
| CD38 | Pacific Blue | 90 | BioLegend | 102720 |
| CD11c | FITC | N418 | BioLegend | 117306 |
| Tbet | BV711 | 4B10 | BioLegend | 644820 |
| Ki-67 | BV605 | 16A8 | BioLegend | 652413 |
| *Antibody-secreting cells panel* | | | | |
| CD19 | BV421 | 6D5 | BioLegend | 115538 |
| B220 | BV785 | RA3-6B2 | BioLegend | 103246 |
| IgM | BV510 | RMM-1 | BioLegend | 406531 |
| IgD | PerCp-Cy5.5 | 11-26c.2a | BioLegend | 405710 |
| IgG1 | BV650 | RMG1-1 | BioLegend | 406629 |
| IgG2b | PE | RMG2b-1 | BioLegend | 406708 |
| CD138 | BV711 | 281-2 | BioLegend | 142519 |
| CD38 | PE-Cy7 | 90 | BioLegend | 102720 |
| Blimp-1 | AF647 | 5E7 | BioLegend | 150004 |

| **Table 1 (continued)** | | | | |
|---|---|---|---|---|
| **Antigen** | **Fluorophore** | **Clone** | **Source** | **Catalog #** |
| *CD4 T cell cytokines panel* | | | | |
| CD4 | Pacific Blue | GK1.5 | BioLegend | 100428 |
| CD8 | PerCp-Cy5.5 | 53-6.7 | BioLegend | 100734 |
| PD-1 | BV711 | 29F.1A12 | BioLegend | 135231 |
| CXCR5 | BV421 | L138D7 | BioLegend | 145511 |
| CD45.1 | AF700 | 104 | BioLegend | 110723 |
| CD45.2 | Spark NIR 685 | A20 | BioLegend | 109863 |
| IL-2 | PE/Cy5 | JES6-5H4 | BioLegend | 503824 |
| IL-4 | PE/Cy7 | 11B11 | BioLegend | 504117 |
| IFN-g | BV650 | XMG1.2 | BioLegend | 505831 |
| IL-17a | AF488 | TX11-18H10.1 | BioLegend | 506909 |
| TNF-a | BV750 | MP6-XT22 | BioLegend | 506358 |
| IL-21 | PE | mhalx21 | Invitrogen | 12-7213-80 |
| *Kidney immunofluorescence* | | | | |
| Anti-mouse | AF647 | | BioLegend | 405322 |
| C3c | FITC | N/A | Invitrogen | PA1-29718 |
| *CD40 signaling antibodies for Immunoblotting* | | | | |
| Akt | N/A | N/A | Cell Signaling | 9272S |
| Phospho-Akt | N/A | N/A | Cell Signaling | 9271T |
| p65 | N/A | D14E12 | Cell Signaling | 8242T |
| Phospho-p65 | N/A | 93H1 | Cell Signaling | 3033T |
| P52/100 | N/A | N/A | Cell Signaling | 4882T |
| B-actin | N/A | 8H10D10 | Cell Signaling | 3700T |
| Rabbit anti- | HRP | N/A | Cell Signaling | 7074S |

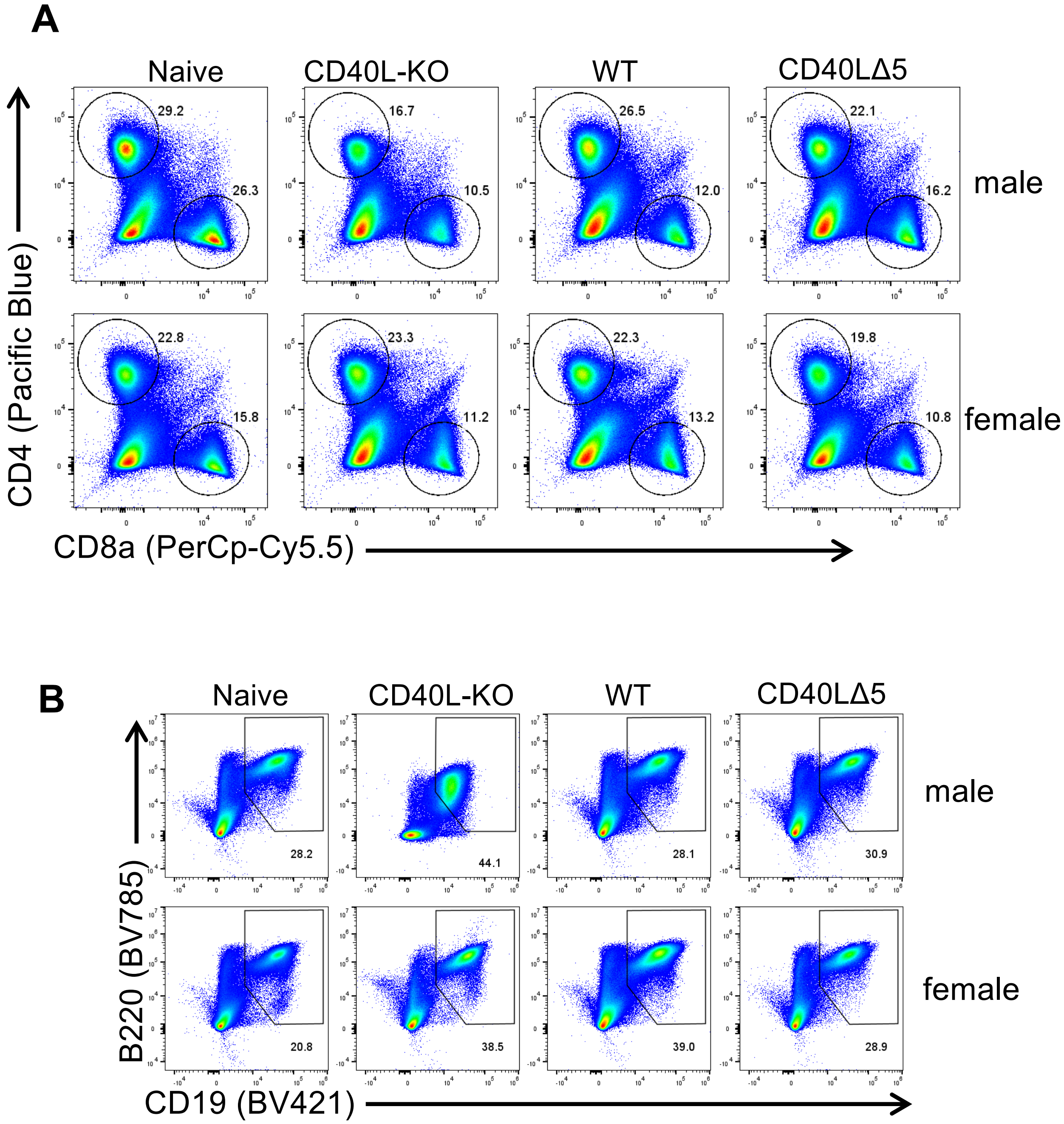


**Supplementary Figure 1.** Evaluation of T and B cell populations in male and female PIL mice showing naïve, CD40L-KO, WT and CD40LΔ5 mice. (A) Spleens were collected 24 weeks post injection with pristane and analyzed for distribution of CD4 and CD8 T cells. (B) The same splenic population was analyzed for numbers and percentages of CD19+CD20+ B cells (bottom).

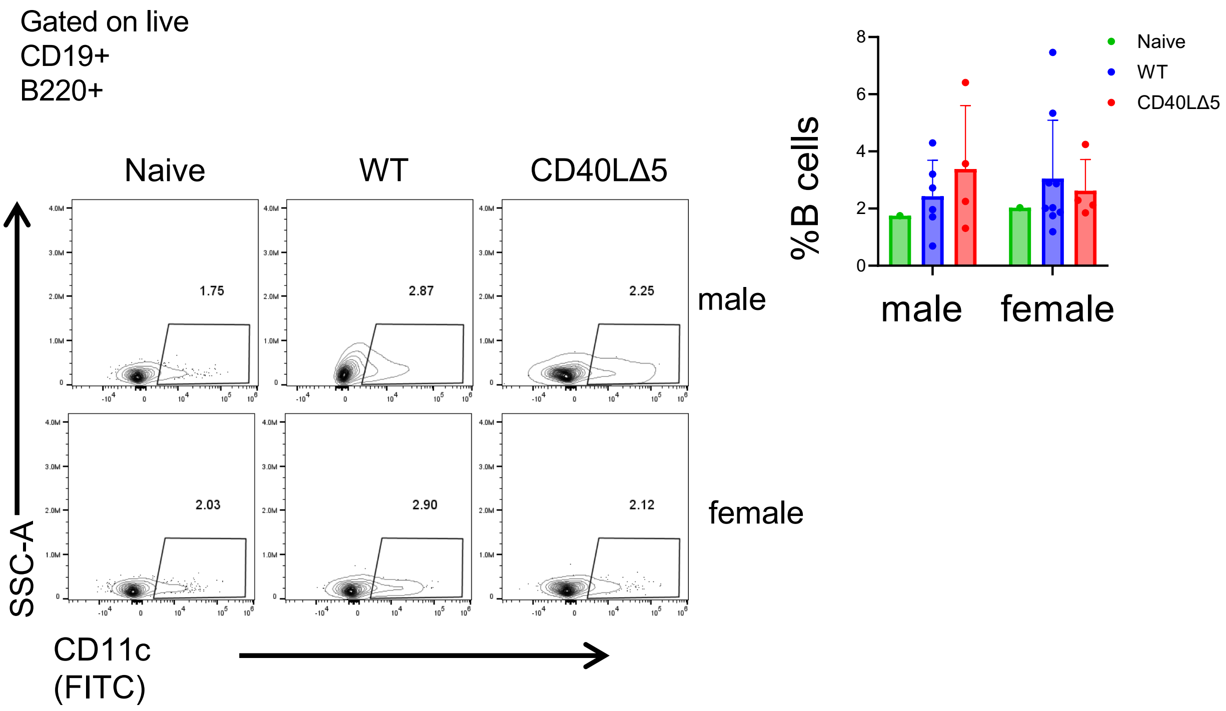


**Supplementary Figure 2.** The CD40LΔ5 mutation does not change the population of Age-associated B cells (ABCs) in PIL mice. Total splenocytes isolated from PIL mice 24 weeks following pristane injection were analyzed for the presence of CD19+CD20+CD11c+ age-associated B cells (ABCs). Representative histograms for male and female naïve, WT, and CD40LΔ5 PIL mice are shown and compiled data is shown in graph form.

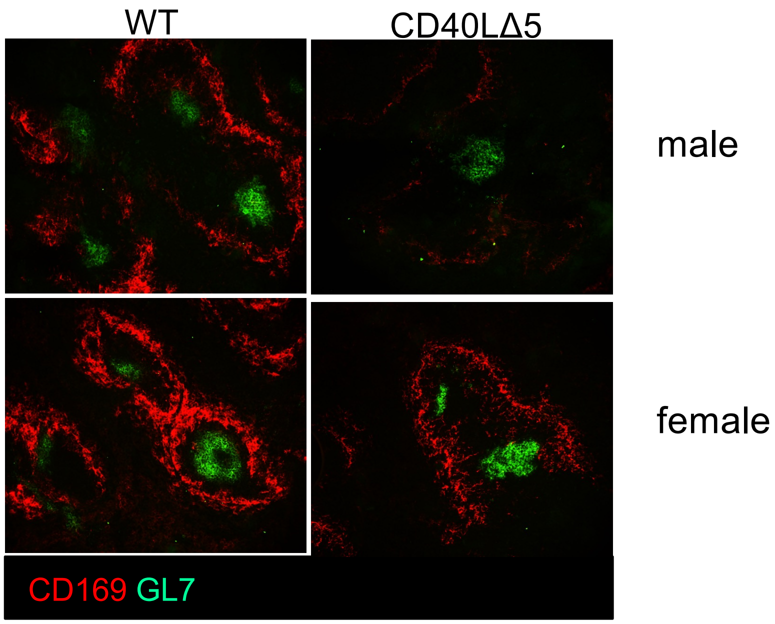


**Supplementary Figure 3.** The effect of the CD40LΔ5 mutation on splenic GC structures in PIL mice. GC structures were visualized in mice 6 months following pristane injection, using 20 mm sections of frozen spleens stained with anti-mouse CD169 to identify follicles (AF594-labeled, red), anti-mouse GL-7 for GC B cells (AF488-labeled, green), and visualized using an Olympus BX63 fluorescent microscope.

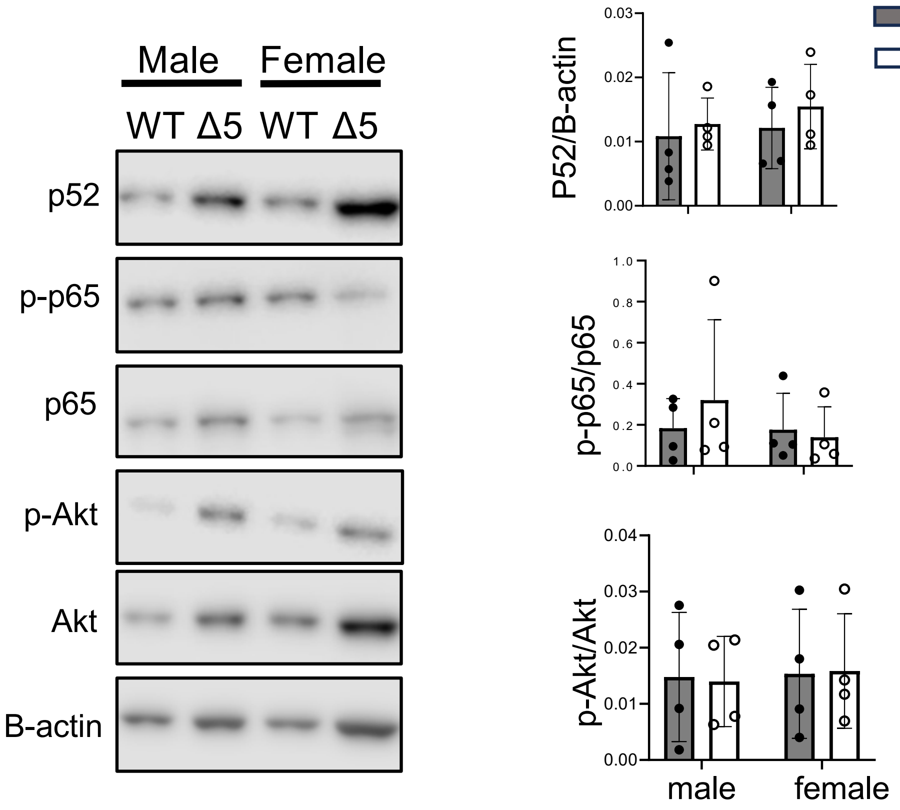


**Supplementary Figure 4.** Activation of CD40 pathways in B cells from WT and CD40LΔ5 mice receiving bm12-WT CD4 T cells or bm12-Δ5 CD4 T cells, respectively. Negative-isolated splenic B cells from 2-wk old recipient mice were analyzed by immunoblot analysis (left). Densitometry analysis was quantified as follows: for p52, values were normalized to those of B-actin; for phosphorylated p65 and Akt, both were normalized to total levels of protein on the same blot. Data are shown as mean ± SEM from two independent experiments
